# Epithelial protein lost in neoplasm (EPLIN)-β is a novel substrate of ornithine decarboxylase antizyme 1, mediating cellular migration

**DOI:** 10.1101/2022.06.16.496394

**Authors:** Dan Li, Suat Peng Neo, Jayantha Gunaratne, Kanaga Sabapathy

**Affiliations:** Division of Cellular & Molecular Research, Humphrey Oei Institute of Cancer Research, National Cancer Centre Singapore, Singapore 169610, Singapore; Institute of Molecular & Cellular Biology, Agency for Science, Technology and Research (A*STAR), Singapore 138673, Singapore; Cancer and Stem Cell Biology Program, Duke-NUS Medical School, Singapore 169857, Singapore; Department of Biochemistry, Yong Loo Lin School of Medicine, National University of Singapore, Singapore 119228, Singapore

**Keywords:** Antizyme, cancer, EPLIN, ODC, LIMA1, Polyamines, ubiquitin-independent degradation, cell migration

## Abstract

Polyamines promote cellular proliferation, and their levels are controlled by ornithine decarboxylase antizyme 1 (Az_1_), through the proteasome-mediated, ubiquitin-independent degradation of ornithine decarboxylase (ODC), the rate-limiting enzyme of polyamine biosynthesis. Az_1_-mediated degradation of other substrates such as cyclin D1, DNp73 and Mps1 also regulate cellular migration and centrosome amplification, and collectively, the currently known six Az_1_-substrates are all linked with tumorigenesis. To understand if Az_1_-mediated protein degradation might play a key role in regulating cellular process associated with tumorigenesis, we employed quantitative proteomics to identify novel Az_1_ substrates. In this report, we describe the identification of LIM domain and Actin-binding protein 1 (LIMA1), also known as epithelial protein lost in neoplasm (EPLIN), as a new target of Az_1_. Interestingly, between the two isoforms of EPLIN (α and β), only EPLIN-β is the substrate of Az_1_, degraded in a proteasome-dependent and ubiquitination-independent manner. Absence of Az_1_ led to elevated EPLIN-β levels, which is causal to enhanced cellular migration of Az_1_^-/-^ cells. Consistently, higher levels of *LIMA1* expression correlated with poorer overall survival of colorectal cancer patients. Taken together, this study identifies EPLIN-β as a novel Az_1_ substrate that regulates cellular migration.

## Introduction

Polyamines, which include putrescine, spermidine, and spermine, are bound to, and modulate the functions of negatively charged molecules such as DNA, RNA and protein (1). They are not only essential for normal cellular growth and differentiation, but also play an important role in cellular proliferation and in the development of cancers (2). Elevated levels of polyamines are associated with multiple cancer types, including breast, colon, lung, prostate, and skin cancers (2), and hence need to be carefully regulated.

Ornithine decarboxylase (ODC) is the rate-limiting enzyme regulating polyamine synthesis (3). This enzyme catalyzes the decarboxylation of ornithine to form putrescine, a common step involved in polyamine biosynthesis leading to the generation of spermidine and spermine. High ODC activity is linked to rapid proliferation of normal and cancerous cells and tissue (4, 5). ODC is negatively regulated by the ornithine decarboxylase antizyme (referred to as Oaz or Az) family, which are encoded by three genes: *OAZ1, OAZ2*, and *OAZ3* in humans (6-8). The most predominant Az protein is Az_1_ which is ubiquitously expressed along with Az_2_, though at higher levels than the latter (6). Az_3_ is however only expressed in the testis (9).

Azs are expressed in a unique manner, as they are derived from two overlapping open reading frames (ORFs): ORF1 contains the translational start codon and also has a stop codon at a frameshift site; and ORF2 encodes most of the protein, and lacks an initiation codon. A +1 ribosomal frameshift results in the skipping of one nucleotide at the stop codon of ORF1 and leads to continued translation to the end of ORF2, which results in the functional full-length Az (10). Increased intracellular polyamine levels stimulate the +1 frameshift, which then negatively regulates ODC expression, as a feedback mechanism to regulate polyamine levels in cells (10-13).

The enzymatically active ODC is a homodimer, and binds to Az with higher affinity to form ODC–Az heterodimers (10). Az binding promotes the degradation of ODC through the 26S proteasome-dependent, independent of ubiquitination (14). Although all Azs are able to inhibit ODC activity and polyamine uptake, only Az_1_ induces ODC degradation (14-16). The role of Az_2_ in ODC degradation is controversial (7, 14, 17), and Az_3_ is unable to induce ODC degradation (8). Therefore, Az_1_-mediated downregulation of ODC and inhibition of polyamine biosynthesis is crucial in the maintenance of polyamine homeostasis.

Az_1_ activity is regulated by an inhibitory interaction with Az inhibitor (AZIN), another member of the broader Az family, which is highly similar to ODC (18, 19). Given the similarity to ODC, AZIN competitively binds Az, thereby relieving ODC from Az-mediated degradation. Thus, the Az/AZIN ratio determines cellular polyamine homeostasis, thereby regulating cellular growth (20). The levels of both Az_1_ and AZIN have been evaluated in several cancers and corresponding normal tissues, which indicate that Az_1_ levels are indeed reduced, with a concomitant increase in AZIN levels in cancers (20), supporting a model for a decrease in Az_1_ levels/activity that promotes cancer development.

Besides ODC, five other proteins whose expression is upregulated in cancers have been identified as Az_1_ substrates thus far, including DNp73, Aurora A, Cyclin D1, Mps1 and Smad1 (21-25). Our laboratory has previously shown that Az_1_ is a negative regulator of DNp73 (25), an oncoprotein of the p53 tumor-suppressor family, that is over-expressed in a variety of cancers and whose expression leads to resistance to a variety of chemotherapeutic drugs and metastasis (26, 27). Az_1_-mediated DNp73 degradation also occurs in an ubiquitin-independent but proteasome-dependent manner, and facilitates chemosensitivity (25), collectively suggesting that Az_1_ could potentially regulate the expression of a larger set of substrates to control tumorigenesis.

In order to uncover the full repertoire of Az_1_ substrates, we undertook a proteomics approach using cells lacking Az_1_ expression, and report here the identification of EPLIN-β as a novel Az_1_ substrate. EPLIN is an actin-binding protein that regulates actin cytoskeleton and its dynamics, and is encoded by the *LIMA1* gene, and is expressed as two isoforms: a 600 amino acid EPLIN-α and a 760 amino acid EPLIN-β, generated from an alternative pre-mRNA splicing events (28). Both isoforms contain a LIM domain, which is a cysteine-rich domain composed of two zinc fingers and functions as modular interface to facilitate protein-protein interactions (29-31). LIM domain containing proteins are frequently present in molecules responsible for cytoskeletal organization (32, 33), and EPLINs also associate with epithelial cell junctions (28, 34-36).

Loss of EPLIN-α expression has been implicated in the progression of various cancers, such as oral, breast, and prostate cancers where its expression was either downregulated or completely abolished compared to normal tissues, indicating a tumor-suppressor role for EPLIN-α (37, 38). On the contrary, increased levels of EPLIN-β have been noted in various cancers (28), suggestive of an oncogenic or growth promoting role for EPLIN-β. Our results show that EPLIN-β, but not EPLIN-α, is degraded in an Az_1_-dependent and ubiquitin-independent manner, although both isoforms were able to interact with Az_1_. Functional analysis indicated that EPLIN-β is a positive regulator of cellular migration, being upregulated and causal to the enhanced cellular growth and migration of Az_1_ null (Az_1_^-/-^) cells. These data therefore demonstrates the identification of EPLIN-β as a novel Az_1_ substrate mediating cellular migration.

## Results

### Generation of Az_1_-deficient cells

In order to identify novel Az_1_ substrates, we generated HCT116 human colorectal cell lacking Az_1_, with the aim of identifying proteins whose expression would be elevated in the absence of Az_1_, using Crispr-Cas9 genome editing. Two pairs of guide (g)RNAs (lentiviral Crispr constructs), targeting different coding regions of *OAZ1* were used (Figure 1A). The first pair of gRNAs targeted ORF1 of *OAZ1*, whereas the second pair targeted ORF2. gRNAs targeting ORF1 resulted in a 684 nucleotide insertion at the 238 nucleotide position, within the Crispr-Cas9 cutting region, and a consecutive 57 nucleotide deletion (corresponding to nucleotides 126-182) in one clone (Figure 1B). The second pair of gRNAs led to the deletion of 92 nucleotides between nucleotides 1963 and 2054 (Figure 1B) in genomic DNA in another clone. Corresponding screening primers and sequencing analysis was performed to confirm these alterations in both clones (Figure 1C). A similar strategy was used to target *OAZ2*, and two representative clones with an insertion of 396 nucleotides and a deletion of 14 nucleotides were obtained (Figure 1B and C).

**Figure 1.**
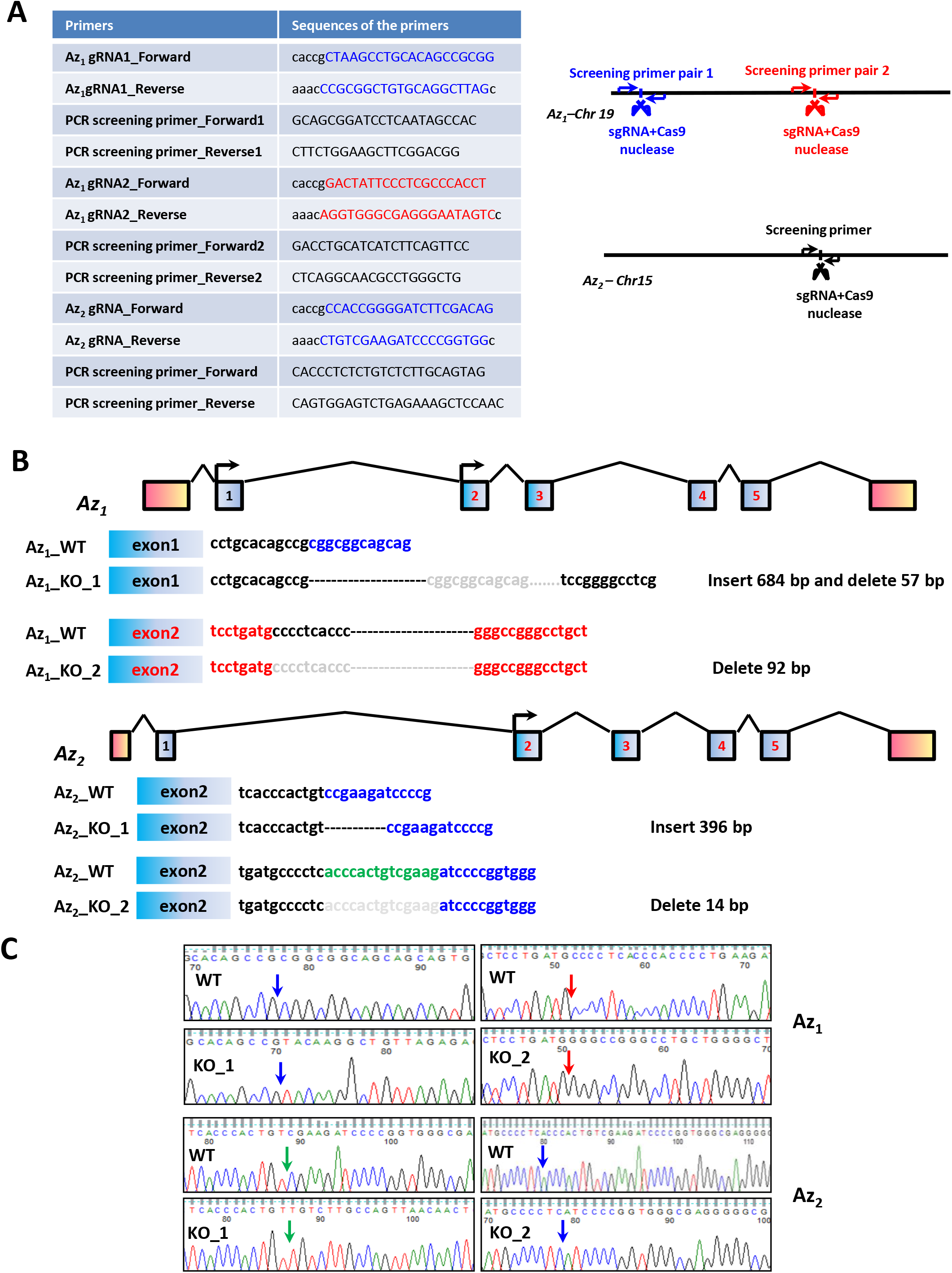
Generation of Az_1_ and Az_2_ knockout cells. (A) Primers used for generating Az_1_ and Az_2_ knockout cells are listed, along with a schematic indicating the positions of the sgRNAs used for targeting. (B) Schematic shows deletions and insertions in two clones each for Az_1_ and Az_2_ used in this study. bp: base pair. (C) Sequencing data for WT and Az_1_^-/-^ and Az_2_^-/-^ HCT116 cells are shown. Arrow indicates the insertion or deletion site on the locus.

### Identification of EPLIN-β as an Az_1_ substrate

Based on the hypothesis that increased levels of Az_1_-substrate proteins could be identified from cells lacking Az_1_, we employed the quantitative SILAC (**S**table **I**sotope **L**abeling using **A**mino acids in **C**ell culture) proteomics methodology (39) and compared the differential expression of proteins between Az_1_^-/-^ cells and their wild-type (WT) controls. Changes in expression of greater than 2-fold (H/L ratios) between heavy and light proteins were considered significant (40). The initial analysis led to the identification of several proteins that were significantly differentially expressed between the Az_1_^-/-^ cells and their WT controls, among which ODC ranked first, validating our experimental approach (Figure 2A). In addition, Cyclin D1 which is another known substrate of Az_1_ was also identified in the analysis, albeit below the H/L ratio of 2, indicating that many more bona-fide substrates could be identified at lower thresholds. Apart from these, several other unique proteins were identified, of which the LIM domain and actin-binding protein 1 *(LIMA1*, which has two isoforms: EPLIN-α and EPLIN-β) was second on the list. Hence, we focused our efforts in characterising the role of Az_1_ in the regulation LIMA1. As aforementioned, EPLIN is expressed as two isoforms, generated from an alternative pre-mRNA splicing event (Figure 2B). Interestingly, our SILAC analysis identified only EPLIN-β as being upregulated in the absence of Az_1_, indicating that the additional sequences in EPLIN-β may be critical for Az_1_-mediated regulation.

**Figure 2.**
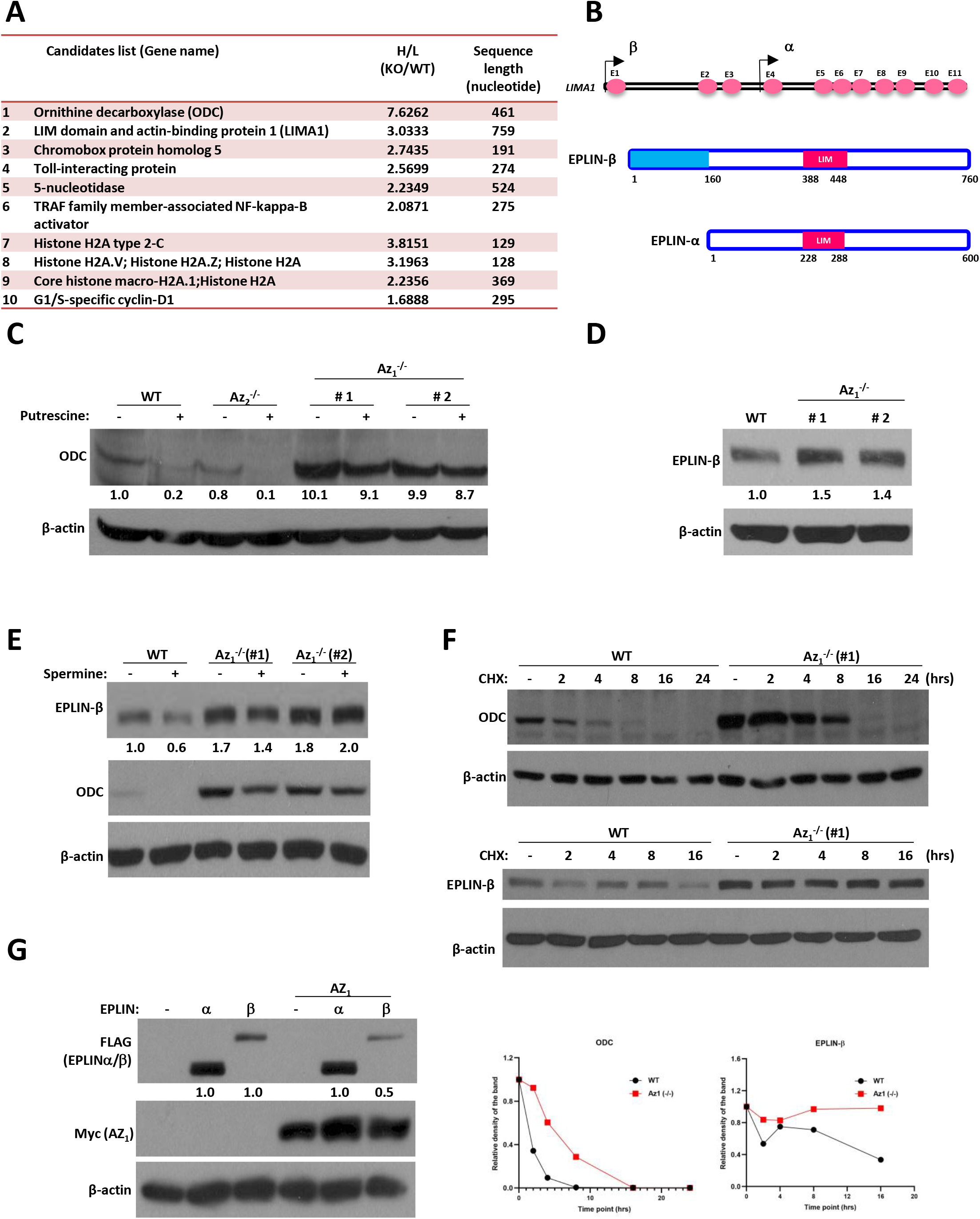
EPLIN-β is a novel substrate of Az_1_. (A) The list shows the top 9 candidates, along with Cyclin D1, identified from the SILAC analysis. (B) Schematic diagram of *LIMA1* gene and structure of the EPLIN-α and -β isoforms. The *LIMA1* gene contains 11 exons and 10 introns. EPLIN-β has an additional 160 amino acids at its amino-terminal which is absent in EPLIN-α. (C-D) HCT116 WT, Az_1_-KO, and Az_2_-KO cells were treated with 2.5 µM putrescine for 24 hrs (C), or were used without any treatment (D). The cells were then harvested and lysed, and immunoblotting was performed to determine the expression of various proteins indicated. Expression levels of ODC (C) and EPLIN-β (D) were quantified relative to β-actin and are indicated below the blots. Expression levels of untreated WT cells were used as a reference (e.g. 1.0). (E-F) WT and Az_1_-KO cells were treated with 50 µM Spermidine for 24 hrs (E), or with 100 µg/ml cycloheximide for the indicated time periods (F), and were harvested and analysed by immunoblotting. (G) H1299 cells were transfected with EPLIN-α and -β plasmids without or with Az_1_ plasmid. 24 hrs post-transfection, the cells were harvested and analyzed by immunoblotting. Multiple blots were run with the same lysates (and same amount) for detection with the various antibodies (e.g. in E and G), and a representative anti-Actin blot for loading control is shown. All experiments were repeated 2-3x independently and representative blots are shown.

To validate if EPLIN-β is indeed regulated by Az_1_, we undertook several approaches. Firstly, we quantified the expression of ODC as well as EPLIN-β levels in Az_1_^-/-^ cells. Baseline ODC levels were low in HCT116 WT cells, but were significantly elevated in two Az_1_^-/-^ clones (Figure 2C), further establishing the utility of the Az_1_^-/-^ cells in the study of its substrates. Absence of Az_2_ did not affect ODC levels, highlighting a dominant role for Az_1_ in ODC regulation, as reported (14). Furthermore, treatment with polyamines (e.g. putrescine), which induces the frameshift of endogenous Az, caused a further decrease in ODC levels only in WT and Az_2_^-/-^ cells, but had no major effects in Az_1_^-/-^ cells (Figure 2C), demonstrating the requirement for Az_1_ in ODC regulation.

Expression of EPLIN-β was also upregulated in the absence of Az_1_ (Figure 2D), but not in the absence of Az_2_ (Supplementary Figure 1A), without any significant impact on *EPLIN-β* mRNA levels (Supplementary Figure 1B). Moreover, while treatment of WT cells with polyamines (e.g. spermine) led to a decrease in EPLIN-β levels, this was not the case in the absence of Az_1_ in which EPLIN-β levels remained relatively unchanged (Figure 2E), further confirming that EPLIN-β levels are indeed regulated in an Az_1_-dependent manner. In addition, treatment of cells with cycloheximide (CHX) to block protein synthesis and subsequent chase indicated that the decay of EPLIN-β and ODC proteins was significantly delayed in the absence of Az_1_ (Figure 2F), suggesting that the effects of Az_1_ on EPLIN-β and ODC levels were at the post-transcriptional level, leading to their extended half-lives.

Finally, we expressed cDNAs encoding either EPLIN-α or EPLIN-β alone or with Az_1_ in H1299 cells to evaluate the direct impact of Az_1_ overexpression on EPLIN levels. Co-expression with Az_1_ led to a decrease in EPLIN-β, but not EPLIN-α levels (Figure 2G). However, co-expression with Az_2_ did not affect EPLIN-β expression (Supplementary Figure 1C). These data collectively establish that in EPLIN-β, but not EPLIN-α, is a target of Az_1_-mediated degradation.

### EPLIN-β interacts with Az_1_ through its LIM domain and is degraded in a ubiquitin-independent manner

As all the reported substrates of Az_1_ interact with it, we next evaluated if EPLIN also interacts with Az_1_. As shown in Figure 3A, both EPLIN-α and EPLIN-β interacted with Az_1_ when co-expressed. Since both EPLIN-α and -β contain a centrally located LIM domain which may mediate self-dimerization or allow EPLIN to interact with other proteins (Figure 3B) (31, 34), we hypothesized that EPLIN may interact with Az_1_ through its LIM domain. To evaluate this possibility, we generated EPLIN-α and -β cDNAs with a LIM domain deletion, which were co-expressed with Az_1._ Absence of LIM domain abrogated binding of both EPLIN isoforms to Az_1_ (Figure 3C), suggesting the presence of additional amino-terminal region in EPLIN-β is critical for its destabilization by Az_1_. Nonetheless, EPLIN-β lacking the LIM domain was refractory to Az_1_-mediated degradation (Figure 3C), emphasizing the importance of interaction between these two proteins for the degradation of EPLIN-β. Moreover, co-expression with AZIN led to partial rescue of EPLIN-β degradation by Az_1_ (Figure 3D), similar to observation with DNp73 and Cyclin D1 (22, 25).

**Figure 3.**
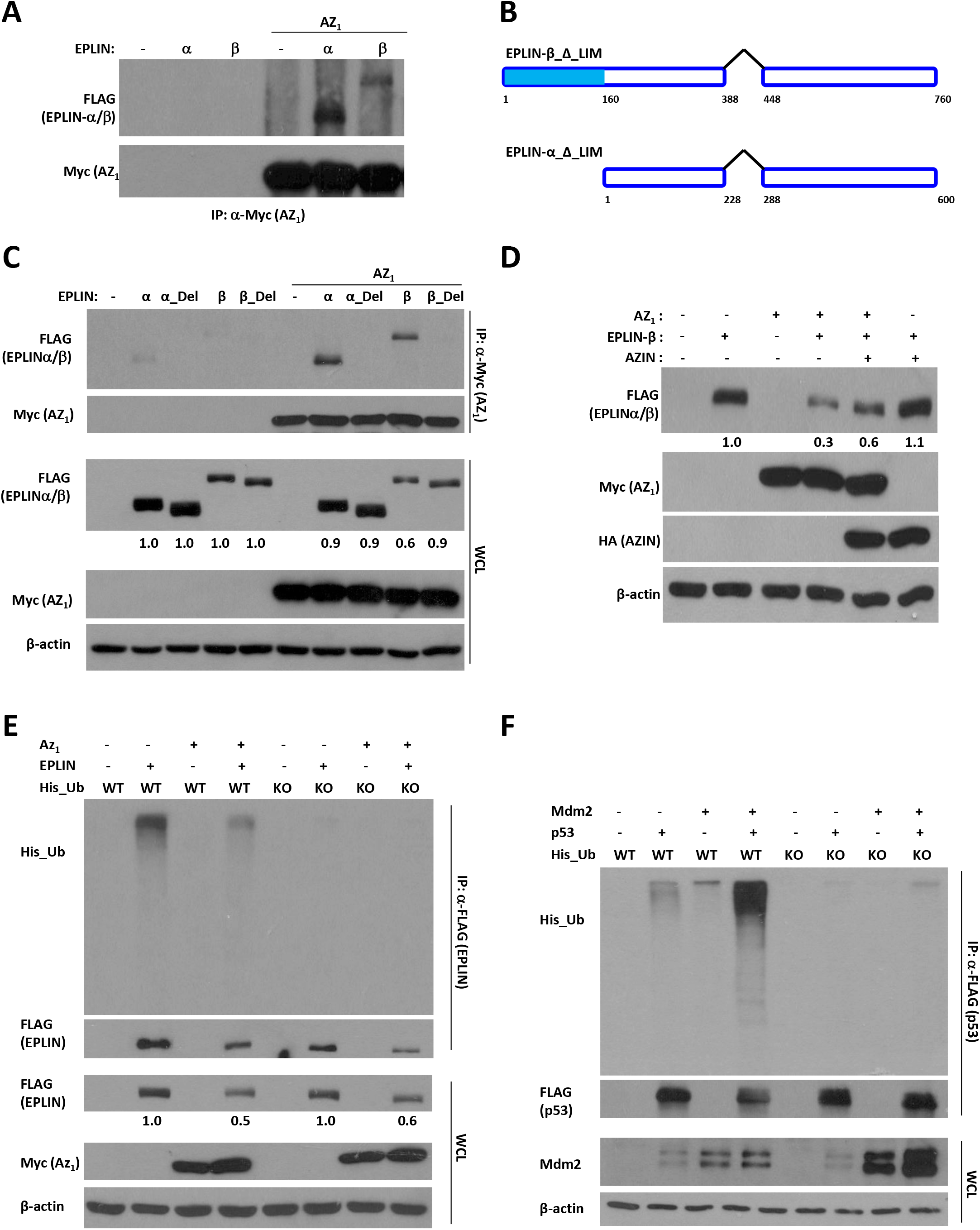
Az_1_ interacts with EPLINs through their LIM domain. (A-D) H1299 cells were transfected with the indicated plasmids, and cellular lysates collected 24 hrs post-transfection were used either for immunoprecipitation (IP) with anti-Myc antibody (A, C), or for direct immunoblotting analysis (D). Schematic structure of the EPLIN-α and -β plasmids used with LIM domain deletion are shown (B). (E-F) H1299 cells were transfected with the indicated plasmids, and used for IP analysis as described above. Multiple blots were run with the same lysates (and same amount) for detection with the various antibodies (e.g. in C-E), and a representative anti-Actin blot for loading control is shown. All experiments were repeated 2-3x independently and representative blots are shown.

We finally evaluated if Az_1_-mediated EPLIN-β degradation occurs independent of ubiquitination, similar to other Az_1_ substrates. To that end, we co-expressed Az_1_ and EPLIN-β along with either WT ubiquitin, or one in which the seven lysine residues required for ubiquitin chain formation have been substituted to arginine residues (i.e. Ubi-KO) (41). EPLIN-β levels were reduced in the presence Az_1_ irrespective of the ubiquitin status (Figure 3E). On the contrary, we used Mdm2-mediated degradation of p53 as a positive control to demonstrate the dependence on ubiquitin chain formation (Figure 3F), as reported (42, 43). These data together demonstrate that EPLIN-β is degraded in an ubiquitin-independent manner by Az_1_ which requires the interaction of these proteins.

### Silencing EPLIN-β expression rescues accelerated cellular growth and migration of Az_1_ ^-/-^ cells

To determine if Az_1_/EPLIN-β interaction has a functional role in cellular growth regulation, we first evaluated the effects of Az_1_-deficiency on cellular growth and migration. Az_1_^-/-^ cells grew much faster than control cells, and led to the formation of a higher number of cellular colonies (Figures 4A). Moreover, absence of Az_1_ led to accelerated cellular migration in would closure assays (area of open wound after 48hrs: control cells *vs*. Az_1_^-/-^ clone 1 *vs*. Az_1_^-/-^ clone 2: 0.38 *vs*. 0.2 *vs*. 0.15 inches) (Figure 4B), together confirming the inhibitory effects of Az_1_ on cellular growth and motility.

**Figure 4.**
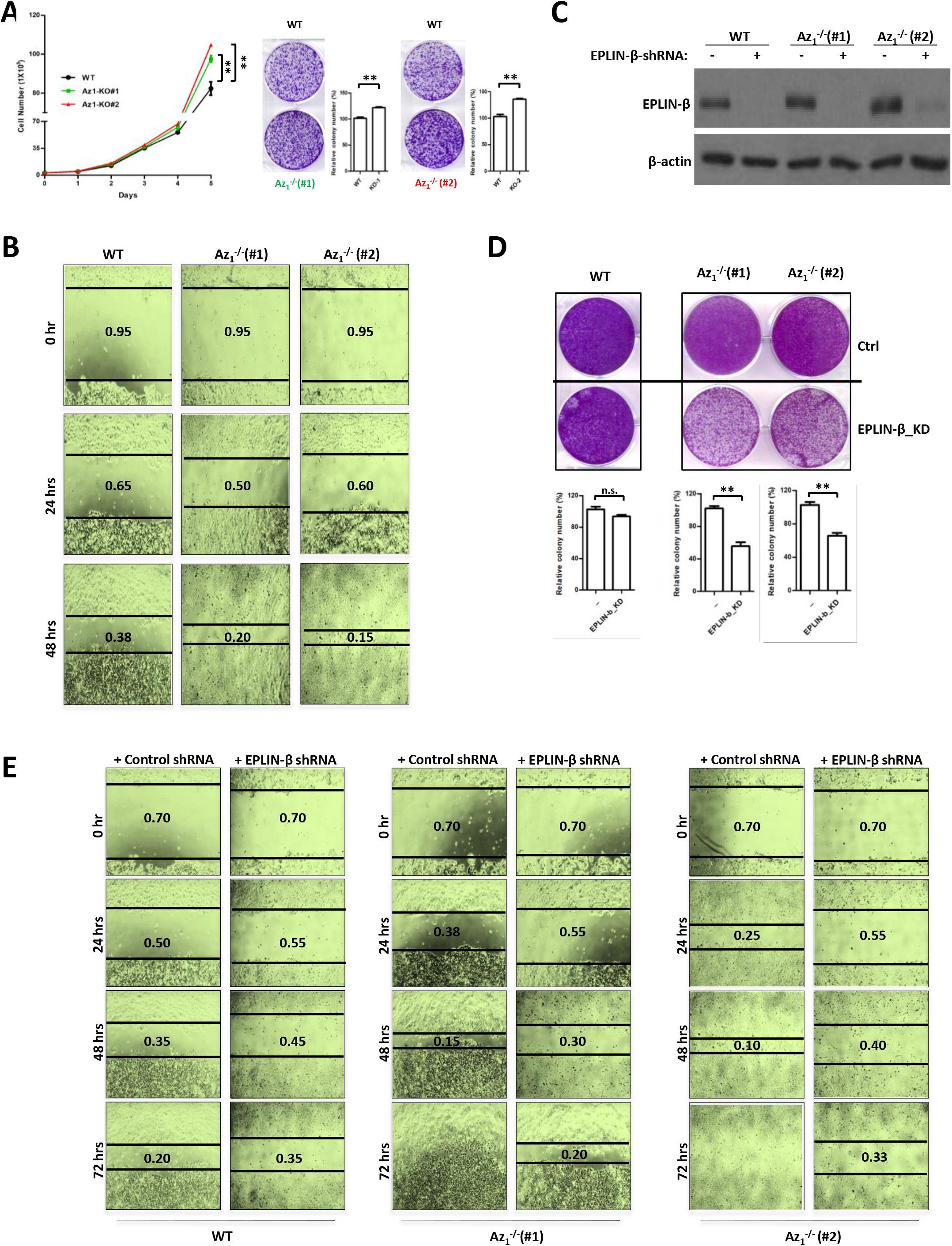
Absence of Az_1_ promotes cell proliferation and migration. (A) Growth rates of two Az_1_-KO clones were compared with WT HCT116 cells over several days (left). X axis represents time (days) after cell seeding. The cells were seeded and grown for 2 weeks prior to crystal-violet (0.5% w/v) staining to determine colony growth (right). (B) Monolayers of WT or Az_1_-KO cells were scraped with a 200-µl sterile pipette tip to generate open wounds, and images were captured at 0, 24 and 48 hrs later (4× magnification) to visualise the wound area that is covered. The distance between the edges were determined and are indicated. (C-E) EPLIN-β expression was silenced using two independent shRNAs in WT and Az_1_-KO cells and the expression levels are shown (C). The cells were used in cellular colony formation (D) and in wound closure assays (E). All experiments were repeated 2-3x independently and representative blots/images are shown.

We next examined if the observed growth advantage could be due to elevated EPLIN-β levels. Although the majority of the previous studies indicate a putative tumor suppressor role for EPLINs, the roles of each individual isoforms (EPLIN-α or EPLIN-β) have not been fully elucidated. Hence, we silenced the expression of EPLIN-β in parental and Az_1_^-/-^ clones (Figure 4C). Silencing EPLIN-β led to a significant reduction in the growth of colonies specifically in both the Az_1_^-/-^ clones, whereas the effect was marginal in the HCT116 WT cells (Figure 4D), indicating that elevated EPLIN-β expression in the absence of Az_1_ indeed contributed to its accelerated growth. Moreover, a similar reversal of the accelerated migration of Az_1_^-/-^ cells was noted in the wound closure assays (area of open wound after 48hrs: control cells *vs*. Az_1_^-/-^ clone 1 *vs*. Az_1_^-/-^ clone 2: 0.35 *vs*. 0.15 *vs*. 0.10 inches, with control shRNA; control cells *vs*. Az_1_^-/-^ clone 1 *vs*. Az_1_^-/-^ clone 2: 0.45 *vs*. 0.3 *vs*. 0.4 inches, with EPLIN-β shRNA) (Figure 4E). These results further illustrate that the increased growth and migration observed in Az_1_^-/-^ cells is due to the upregulation of EPLIN-β.

### High levels of EPLIN-β prognosticate poorer survival rates

The above data suggests that EPLIN-β may function as a cellular growth promoter rather than a tumor suppressor. To assess EPLIN-β’s expression in cancers, we evaluated and found that *LIMA1* (encoding EPLIN-β) expression was higher in tumor tissues compared to normal tissues in colorectal cancers, based on the UALCAN website (Figure 5A). Furthermore, Kaplan-Meier analysis of overall survival in different colorectal carcinoma datasets indicated that higher levels of *LIMA1* (EPLIN-β) correlated with a significantly lower survival probability (Figure 5B, left panel). Interestingly, higher *OAZ1* levels correlated with a higher survival probability in the same datasets (Figure 5B, right panel), albeit with much lower significance. These data suggest that EPLIN-β which enhances cancer cell growth and migration, is indeed associated with poorer patient survival.

**Figure 5.**
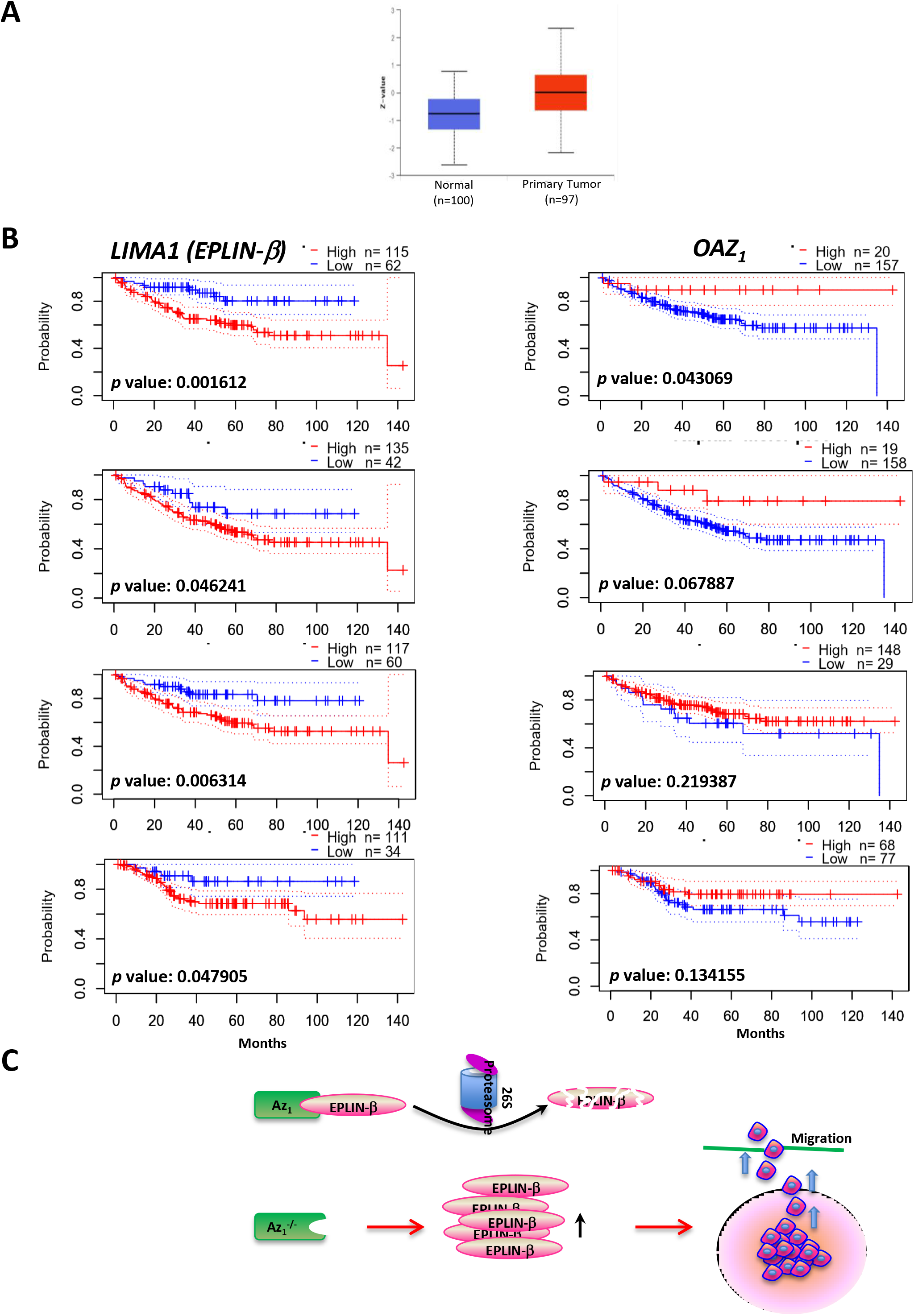
LIMA1 or OAZ1 expression and prognosis among colorectal cancer patients. (A) Higher *LIMA1* gene expression was noted in primary tumor tissues compared to normal tissues from colorectal cancer (CRC) patients, based on the analysis of the UALCAN database (http://ualcan.path.uab.edu/index.html). (B) Correlation between *LIMA1* (EPLIN-β) (left) and *OAZ1* (right) expression with CRC patients’ prognosis was determined using PrognoScan, and Kaplan-Meier plots are shown. The same datasets were used in both cases. Patients were segregated based on high (red) or low (blue) expression levels. The number of patients in each group is shown at the top of each graph. (C) Working model: EPLIN-β is a novel substrate of Az_1_. The interaction of EPLIN-β with Az_1_ results in its degradation in an ubiquitination-independent and proteasome-dependent manner. Absence of Az_1_ leads to increased EPLIN-β levels, which promote the migration of HCT116 cells.

## Discussion

This report describes the identification of EPLIN-β as a novel Az_1_ substrate. This work expands the repertoire of Az_1_ substrates to seven, and also suggests the existence of many more substrates which are yet to be identified and characterized. Our proteomics analysis identified around 10 substrates in HCT116 colorectal cells which were upregulated significantly (at H/L>2) in the absence of Az_1_, of which we have characterized EPLIN-β here, and the others are being currently investigated. We envisage that other substrates that are regulated in a cell type and context-dependent manner are likely to exist, and are being explored in our laboratory. The growing list of Az_1_ substrates prompts us to speculate that Az_1_–mediated protein degradation might be a yet underexplored regulatory mechanism that could be critical in the regulation of cellular growth, and thus in cancer development, as well as in other physiological and pathological processes.

Polyamines are ubiquitously produced, and their levels are relatively high in rapidly growing cells, and in particular, increased polyamines levels have been shown to directly correlate with disease activity and tumor burden (44, 45). Polyamines lead to the synthesis of Azs, which then negatively regulates polyamine production through the binding and proteasome-mediated degradation of ODC, the rate-limiting enzyme involved in polyamine biosynthesis (46).

ODC, being the classical bona-fide substrate of Az_1_, has been utilized to study the mechanistic basis of Az_1_-mediated degradation. Binding of Az_1_ to ODC leads to disruption of ODC homodimers, leading to the exposure of the C-terminal tail, which targets ODC to the proteasome, in an ubiquitin-independent manner (3, 15, 47, 48). For a long time, no other substrates were identified, but five other substrates, such as Mps1, Smad1, Cyclin D1, Aurora A and DNp73, were later found to be degraded in an Az_1_–dependent manner. All of them have been shown to bind to Az_1_ and are targeted for degradation in an ubiquitin-independent manner (21-25).

Interestingly, a few features appear to be common among all the seven substrates identified thus far: they act as homodimers, or can heterodimerize with highly identical isoform variants; and they are all involved in promoting cellular proliferation and/or migration, being overexpressed in a variety of cancers (49-53). All seven substrates of Az_1_ including EPLIN-β are negatively regulated by Az_1_, consistent with the latter’s growth inhibitory properties. Nevertheless, whether Az_1_–dependent degradation is indeed a relevant mechanism of proteolysis is unclear, due to the lack of further insights into its substrates. Our work was initiated to explore the Az_1_–substratome, using Az_1_^-/-^ HCT116 colorectal cells. Absence of Az_1_ expression expectedly led to elevated levels of ODC, which was also identified as the top candidate in the SILAC analysis. This proteomics analysis also identified a list of proteins that are differentially expressed in the absence of Az_1_, suggesting that the Az_1_–substratome might be much larger than expected, and is the subject of our further investigations.

Similar to the other Az_1_ substrates, EPLIN-β was able to interact with Az_1_ and was degraded in an Az_1_–dependent manner, without the need for ubiquitination. Co-expression with Az_1,_ or treatment with polyamines was sufficient for EPLIN-β degradation in an Az_1_– dependent manner. Furthermore, the enhanced cellular growth and migration phenotypes observed due to Az_1_ absence was reversed by silencing EPLIN-β expression, demonstrating the functional interaction between Az_1_ and EPLIN-β. Consistently, we also found that high EPLIN-β levels or lower Az_1_ levels prognosticated for poorer survival, further confirming the functionality of the data in the clinical context.

EPLIN-β is an actin-binding protein, and its negative regulation by Az_1_ extends a role for latter in cellular migration, expanding its repertoire of tumor suppressive functions. Az_1_ has been suggested to be a tumor suppressor, with its expression being reduced in many cancer types (35). Previous analysis of cyclin D1 as an Az_1_ substrate has suggested a role for it in cellular proliferation/growth (22). Consistently, Az_1_^-/-^ cells had a growth advantage, and we also noticed that they migrated faster in cellular wounding assays compared to their WT counterparts. Furthermore, silencing of EPLIN-β reversed the enhanced migration of Az_1_^-/-^ cells, further demonstrating an important role for Az_1_ in regulating cellular motility.

Among the two EPLIN isoforms, EPLIN-α expression is significantly downregulated in several cancers (compared to normal tissues), leading to enhanced migration or invasion capabilities (28). Overexpression of EPLIN-α leads to inhibition of cellular growth (54, 55). Hence, EPLINs were generally considered as tumor suppressors. However, it is to be noted that cancers also exhibit a concomitant increase in EPLIN-β expression (28), though it has not been clarified whether the reduction in EPLIN-α or the increase in EPLIN-β, or a combination of both, is causal to the enhanced cellular migration. Our results indicate that, at least in HCT116 colorectal cancer cells, EPLIN-β plays a pro-growth role, as opposed to EPLIN-α’s established tumor suppressor functions (54, 55). Consistently, analysis of several human colorectal cancer gene expression datasets showed that survival rates was significantly reduced among the patients whose tumors expressed relatively higher levels of *LIMA1* (denoting EPLIN-β) compared with those expressing lower *LIMA1* levels. This further indicates that EPLIN-α expression is either downregulated or undetectable in colorectal cancers, with EPLIN-β being upregulated, consistent with previous observations (28).

Finally, several points are worth highlighting, based on our results. Firstly, as with other substrates, binding of Az_1_ alone is insufficient, although it is necessary for EPLIN-β degradation. As such, the other EPLIN-α isoform, which is also able to bind to Az_1_, is not degraded by Az_1_. This observation is similar to the p73 proteins, of which DNp73, but not the TAp73 isoform, is degraded by Az_1_, albeit both being able to bind Az_1_ (25). This indicates that the presence of additional domains is required for the degradation process. Next, Az_1_, but not Az_2_, was the key mediator of EPLIN-β degradation. Overexpression of Az_2_ or its absence did not alter EPLIN-β levels. Thus, although polyamines can induce the frame-shifting of both Az_1_ and Az_2_, only the former appears to be the major regulator of protein abundance, at least in the cellular systems we have tested. Thirdly, it is to be noted that many of the newly identified substrates have a much lower H/L ratio compared to ODC. This probably provides an explanation as to why other Az_1_-substrates have not been identified thus far. Moreover, we have detected the interaction between Az_1_ and EPLIN-β using overexpression systems, as it is technically difficult to detect interaction of endogenous proteins as antibodies against processed full-length Az_1_ are currently not available commercially.

Taken together, the data presented in this report provides evidence that EPLIN-β is a novel substrate of Az_1_, mediating Az_1_-dependent cellular growth and migration (Figure 5C). This study also highlights the existence of other substrates of Az_1_, warranting the further exploration of the entire Az_1_-substrome, which could shed light on potential new targets that could be useful in clinical investigations. Moreover, this study also suggests that activation of Az_1_ could be a strategy to inhibit the growth and migration of tumor cells.

## Materials and Methods

### Cell culture and transfection

p53-null human lung cancer H1299 cells, and p53-proficient human colorectal carcinoma HCT116 cells were used in this study. Cells were grown in DMEM supplemented with 10% bovine fetal serum (FBS; Hyclone), 1% penicillin–streptomycin solution, 2 mM L-glutamine (Invitrogen, Carlsbad, CA, USA), 100 μM non-essential amino acids (Invitrogen) and 0.1 mM sodium pyruvate (Invitrogen), as described (56). The EPLIN-β knock-down cells were maintained in complete DMEM medium containing 0.5 µg/ml puromycin.

The relevant plasmids were transfected using Lipofectamine 2000 in accordance with the manufacturer’s instructions (Invitrogen). Expression vectors (all in pcDNA3.0) expressing full length EPLIN-α/-β, Az_1_, ODC, or AZIN have been described previously (57, 58).

### <u>S</u>table <u>I</u>sotope <u>L</u>abeling using <u>A</u>mino acids in <u>C</u>ell culture (SILAC)

HCT116 parental cells were cultured with ‘Heavy’ or ‘Light’ medium and the corresponding Az_1_-KO cells were cultured with ‘Light’ or ‘Heavy’ medium for 4∼5 doublings, at a cell-density of around 20-30% confluence. The cells were harvested, and normalized protein extracts were used for SILAC analysis. LDS buffer (Invitrogen) and reducing agent (Invitrogen) were added to the lysates and boiled for 5 minutes. The proteins were then separated on 4–12% NuPage Novex Bis–Tris Gel (Invitrogen); stained with colloidal blue staining kit (Invitrogen) and digested with trypsin using in-gel digestion procedures (59).

### Mass spectrometry and data analysis

Tryptic peptides were analysed using an EASY-nLC 1000 coupled to a Q Exactive™ Hybrid Quadrupole-Orbitrap (Thermo Fisher Scientific). The peptides were resolved and separated on a 50 cm analytical EASY-Spray column equipped with pre-column over a 120 min gradient ranging from 8 to 38% of 0.1% formic acid in 95% acetonitrile/water at a flow rate of 200nl/min. Survey full scan MS spectra (m/z 310–2000) were acquired with a resolution of 70k, an AGC target of 3 × e^6^ and a maximum injection time of 10 ms. Top twenty most intense peptide ions in each survey scan were sequentially isolated to an ACG target value of 5 x e^4^ with resolution of 17,500 and fragmented using normalized collision energy of 25. A dynamic exclusion of 10s and isolation width of 2 m/z were applied. SILAC peptide and protein quantification was performed with MaxQuant version 1.5.0.30 using default settings. Database searches of MS data were performed using Uniprot human fasta (2017) with tryptic specificity allowing maximum two missed cleavages, two labeled amino acids, and an initial mass tolerance of 4.5 ppm for precursor ions and 0.5 Da for fragment ions. Cysteine carbamidomethylation was searched as a fixed modification, and N-acetylation and oxidized methionine were searched as variable modifications. Labeled arginine and lysine were specified as fixed modifications. Maximum false discovery rates were set to 0.01 for both protein and peptide. Proteins were considered identified when supported by at least one unique peptide with a minimum length of seven amino acids.

### Immunoblot and immunoprecipitation analysis

Cell lysates were prepared in lysis buffer containing 0.5% Nonidet P-40 (NP-40) with protease inhibitor cocktail and dithiothreitol as described (58). The total protein were quantified and then boiled in 4×SDS sample buffer followed by separation on SDS polyacrylamide gels. Immunoblotting was performed with the antibodies, as follows: anti-actin (Sigma), and anti-Flag antibody (Thermo Fisher Scientific Inc., Austin, TX, United States); anti-EPLIN and anti-Myc tag antibodies (Santa Cruz Biotechnology Inc., Dallas, TX, United States); anti-mouse secondary and anti-rabbit secondary antibodies, anti-HA tag and anti-His tag antibodies (Cell Signaling Technology Inc., Danvers, MA, United States).

Cell lysates, prepared as afore-described, were used for co-immunoprecipitation (IP) assays as described previously (57). Briefly, cell lysates were immunoprecipitated with the indicated antibodies for 3 h at 4 °C and then incubated with protein G (GE Healthcare Life Sciences) for 2 hrs at 4 °C. Protein G-agarose was washed with NP-40 buffer three times and then analyzed by western blot with the indicated antibodies.

### RNA-guided CRISPR-Cas9 nuclease-mediated Az knock-out

To knockout Az_1_ or Az_2_ in HCT116 cells, Crispr-Cas9 system was used as previously published (60). Briefly, sgRNAs targeting Az_1_ or Az_2_ were designed using the online tool (http://crispor.tefor.net/) and cloned into the vectors lentiCRISPRv2 (Addgene plasmid # 52961) using Bbs1 enzyme. After validation by DNA sequencing, the sgRNA vectors were co-transfected with corresponding packaging plasmids into 293T cells, and the virus supernatant was prepared. HCT116 cells were then transduced with polybrene (4 mg/ml, Sigma) and viral supernatant, and selected on 0.5μg/ml puromycin containing media for 10-12 days. Single colonies were randomly picked up into 96-well plates for genotyping by PCR and the knockouts were further validated by PCR using the Az_1_ or Az_2_ specific genomic primers (Figure 1).

### Protein half-life and ubiquitination assays

To determine EPLIN-β half-life, HCT parental and Az_1_-KO cells were seeded into 6-well plate at ∼60% confluence. After 24 hrs, the cells were treated with the protein synthesis inhibitor cycloheximide (Sigma) for the indicated durations before harvesting and analysis by immunoblotting.

For EPLIN-β ubiquitination analysis, H1299 cells were transfected with His-ubiquitin, Myc-Az_1_, or FLAG-EPLIN-β. 24 hrs post-transfection, cells were lysed in NP-40 lysis buffer and then incubated with anti-FLAG antibody for 3 h and protein G-agarose beads for 2 hrs at 4 °C. After washing three times, the ubiquitinated EPLIN-β was detected by Immunoblotting using anti-His monoclonal antibody.

### RNA extraction and real-time RT-PCR

Total RNA was prepared from cells using TRIzol reagent (Invitrogen) according to the manufacturer’s instructions. 1.5–2 µg of total RNA was reverse transcribed into cDNA using SuperscriptII (Invitrogen). The sequences of real-time PCR primers and semi-quantitative reverse transcription-PCR analysis are the same as described (56).

### Cellular colony formation and scratch-wound healing (migration) assays

HCT116 cells - WT or Az_1_ knockout, or Az_1_ knockout cells with or without EPLIN-β knock-down cells (1000 cells per well), were plated on 6-well plates and grown for two weeks. After rinsing three times with PBS, cells were fixed with 4% paraformaldehyde in PBS, and stained with 1% crystal violet.

Similarly, the above cells were seeded in 24-well plates at a density of 1.6×10^4^ cells/well in complete DMEM medium and cultured to confluence. The cells were then serum-starved overnight in DMEM medium, and the confluent cell monolayers were then scraped with a yellow pipette tip to generate scratch wounds, and were washed twice with Opti-MEM medium to remove cell debris (61). Cells were incubated at 37°C with the medium containing 0.5 µg/ml puromycin. Time-lapse images were captured at 0, 24, 48 and 72 hr time points in the same position using a Nikon Eclipse TE2000-5 microscope.

### Statistical analysis

Data were analysed by two-way analysis of variance. The differences in mean values were considered significant at *p* values ≤ 0.01(**), and ≤ 0.05(*). The log-rank test was applied in the Kaplan–Meier survival examination. Statistical comparisons between two groups were carried out by Student’s *t*-test and one-way analysis of variance. A two-tailed *p*-value <0.05 was considered significant.

## Acknowledgements

We thank Catherine Kok for technical assistance. This research was supported by the: National Medical Research Council Singapore; NCC Cancer Fund (all to KS); and NCC Cancer Fund (NCCSCF-R-YR2021-OCT-DP1) to DL.

## Author contributions

DL and KS conceived the idea of evaluating the Az_1_-substratome and planned the experiments. DL performed all experiments. SPN and JG supported with all the SILAC-proteomics analysis. DL and KS wrote the manuscript with input from all others. All authors reviewed and commented on the manuscript.

## Supplementary Information

### Supplementary Figure Legends

***Supplementary Figure 1. Analysis of the effects of Az***_***1***_ ***and Az***_***2***_ ***on Eplin-β***

(A-B) HCT116 parental (WT) and Az_2_-KO cells (A) or Az_1_-KO cell clones (B) were harvested and used for immunoblotting analysis (A), or the RNA was used to determine *EPLIN-β* by real-time qPCR analysis (B).

(C) H1299 cells were transfected with the indicated plasmids, and harvested 24 hrs post-transfection and used for immunoblotting analysis.

***Supplementary Table 1. Primer list***

Primers for cloning the indicated plasmids are listed.

## References

1. Igarashi K & Kashiwagi K (2000) Polyamines: mysterious modulators of cellular functions. Biochemical and biophysical research communications 271(3):559–564.

2. Nowotarski SL, Woster PM, & Casero RA (2013) Polyamines and cancer: implications for chemotherapy and chemoprevention. Expert reviews in molecular medicine 15.

3. Palanimurugan R, Scheel H, Hofmann K, & Jürgen Dohmen R (2004) Polyamines regulate their synthesis by inducing expression and blocking degradation of ODC antizyme. The EMBO journal 23(24):4857–4867.

4. Porter CW, Herrera‐Ornelas L, Pera P, Petrelli NF, & Mittelman A (1987) Polyamine biosynthetic activity in normal and neoplastic human colorectal tissues. Cancer 60(6):1275–1281.

5. Gerner EW & Meyskens FL (2004) Polyamines and cancer: old molecules, new understanding. Nature Reviews Cancer 4(10):781–792.

6. Ivanov IP, Gesteland RF, & Atkins JF (1998) A second mammalian antizyme: conservation of programmed ribosomal frameshifting. Genomics 52(2):119–129.

7. Chen H, MacDonald A, & Coffino P (2002) Structural elements of antizymes 1 and 2 are required for proteasomal degradation of ornithine decarboxylase. Journal of Biological Chemistry 277(48):45957–45961.

8. Snapir Z, Keren-Paz A, Bercovich Z, & Kahana C (2009) Antizyme 3 inhibits polyamine uptake and ornithine decarboxylase (ODC) activity, but does not stimulate ODC degradation. Biochemical Journal 419(1):99–104.

9. Ivanov IP, Rohrwasser A, Terreros DA, Gesteland RF, & Atkins JF (2000) Discovery of a spermatogenesis stage-specific ornithine decarboxylase antizyme: antizyme 3. Proceedings of the National Academy of Sciences 97(9):4808–4813.

10. Matsufuji S, et al. (1995) Autoregulatory frameshifting in decoding mammalian ornithine decarboxylase antizyme. Cell 80(1):51–60.

11. Rom E & Kahana C (1994) Polyamines regulate the expression of ornithine decarboxylase antizyme in vitro by inducing ribosomal frame-shifting. Proceedings of the National Academy of Sciences 91(9):3959–3963.

12. Agostinelli E, et al. (2010) Polyamines: fundamental characters in chemistry and biology. Amino acids 38(2):393–403.

13. Coffino P (2001) Regulation of cellular polyamines by antizyme. Nature reviews Molecular cell biology 2(3):188–194.

14. Murakami Y, et al. (1992) Ornithine decarboxylase is degraded by the 26S proteasome without ubiquitination. nature 360(6404):597–599.

15. Li X & Coffino P (1993) Degradation of ornithine decarboxylase: exposure of the C-terminal target by a polyamine-inducible inhibitory protein. Molecular and cellular biology 13(4):2377–2383.

16. Mangold U & Leberer E (2005) Regulation of all members of the antizyme family by antizyme inhibitor. Biochemical Journal 385(1):21–28.

17. Zhu C, Lang DW, & Coffino P (1999) Antizyme2 is a negative regulator of ornithine decarboxylase and polyamine transport. Journal of Biological Chemistry 274(37):26425–26430.

18. Qiu S, Liu J, & Xing F (2017) Antizyme inhibitor 1: a potential carcinogenic molecule. Cancer Science 108(2):163–169.

19. Nilsson J, Grahn B, & Heby O (2000) Antizyme inhibitor is rapidly induced in growth-stimulated mouse fibroblasts and releases ornithine decarboxylase from antizyme suppression. Biochemical Journal 346(3):699–704.

20. Olsen RR & Zetter BR (2011) Evidence of a role for antizyme and antizyme inhibitor as regulators of human cancer. Molecular Cancer Research 9(10):1285–1293.

21. Lim S & Gopalan G (2007) Antizyme1 mediates AURKAIP1-dependent degradation of Aurora-A. Oncogene 26(46):6593–6603.

22. Newman RM, et al. (2004) Antizyme targets cyclin D1 for degradation: A novel mechanism for cell growth repression. Journal of Biological Chemistry 279(40):41504–41511.

23. Kasbek C, Yang C-H, & Fisk HA (2010) Antizyme restrains centrosome amplification by regulating the accumulation of Mps1 at centrosomes. Molecular biology of the cell 21(22):3878–3889.

24. Gruendler C, Lin Y, Farley J, & Wang T (2001) Proteasomal degradation of Smad1 induced by bone morphogenetic proteins. Journal of Biological Chemistry 276(49):46533–46543.

25. Dulloo I, Gopalan G, Melino G, & Sabapathy K (2010) The antiapoptotic DeltaNp73 is degraded in a c-Jun–dependent manner upon genotoxic stress through the antizyme-mediated pathway. Proceedings of the National Academy of Sciences 107(11):4902–4907.

26. Müller M, et al. (2005) TAp73/ΔNp73 influences apoptotic response, chemosensitivity and prognosis in hepatocellular carcinoma. Cell Death & Differentiation 12(12):1564–1577.

27. Steder M, et al. (2013) DNp73 exerts function in metastasis initiation by disconnecting the inhibitory role of EPLIN on IGF1R-AKT/STAT3 signaling. Cancer cell 24(4):512–527.

28. Maul RS & Chang DD (1999) EPLIN, epithelial protein lost in neoplasm. Oncogene 18(54):7838–7841.

29. Kang S, et al. (2000) PCD1, a novel gene containing PDZ and LIM domains, is overexpressed in several human cancers. Cancer research 60(18):5296–5302.

30. Sánchez-García I & Rabbits TH (1994) The LIM domain: a new structural motif found in zinc-finger-like proteins. Trends in Genetics 10(9):315–320.

31. Schmeichel KL & Beckerle MC (1994) The LIM domain is a modular protein-binding interface. Cell 79(2):211–219.

32. Brown MC, Perrotta JA, & Turner CE (1996) Identification of LIM3 as the principal determinant of paxillin focal adhesion localization and characterization of a novel motif on paxillin directing vinculin and focal adhesion kinase binding. The Journal of cell biology 135(4):1109–1123.

33. Collins RJ, Jiang WG, Hargest R, Mason MD, & Sanders AJ (2015) EPLIN: a fundamental actin regulator in cancer metastasis? Cancer and Metastasis Reviews 34(4):753–764.

34. Maul RS, et al. (2003) EPLIN regulates actin dynamics by cross-linking and stabilizing filaments. The Journal of cell biology 160(3):399–407.

35. Abe K & Takeichi M (2008) EPLIN mediates linkage of the cadherin–catenin complex to F-actin and stabilizes the circumferential actin belt. Proceedings of the National Academy of Sciences 105(1):13–19.

36. Song Y, Maul RS, Gerbin CS, & Chang DD (2002) Inhibition of anchorage-independent growth of transformed NIH3T3 cells by epithelial protein lost in neoplasm (EPLIN) requires localization of EPLIN to actin cytoskeleton. Molecular biology of the cell 13(4):1408–1416.

37. Liu Y, Sanders AJ, Zhang L, & Jiang WG (2012) EPLIN-α expression in human oesophageal cancer and its impact on cellular aggressiveness and clinical outcome. Anticancer research 32(4):1283–1289.

38. Liu R, et al. (2016) Epithelial protein lost in neoplasm-α (EPLIN-α) is a potential prognostic marker for the progression of epithelial ovarian cancer. International journal of oncology 48(6):2488–2496.

39. Ong S-E, et al. (2002) Stable isotope labeling by amino acids in cell culture, SILAC, as a simple and accurate approach to expression proteomics. Molecular & cellular proteomics 1(5):376–386.

40. Duan X, Kelsen SG, Clarkson Jr AB, Ji R, & Merali S (2010) SILAC analysis of oxidative stress‐ mediated proteins in human pneumocytes: New role for treacle. Proteomics 10(11):2165–2174.

41. Stringer DK & Piper RC (2011) Terminating protein ubiquitination: Hasta la vista, ubiquitin. Cell Cycle 10(18):3067–3071.

42. Oliner JD, et al. (1993) Oncoprotein MDM2 conceals the activation domain of tumour suppressor p53. Nature 362(6423):857–860.

43. Haupt Y, Maya R, Kazaz A, & Oren M (1997) Mdm2 promotes the rapid degradation of p53. Nature 387(6630):296–299.

44. Soda K (2011) The mechanisms by which polyamines accelerate tumor spread. Journal of Experimental & Clinical Cancer Research 30(1):1–9.

45. Durie BG, Salmon SE, & Russell DH (1977) Polyamines as markers of response and disease activity in cancer chemotherapy. Cancer research 37(1):214–221.

46. Kahana C (2018) The antizyme family for regulating polyamines. Journal of Biological Chemistry 293(48):18730–18735.

47. Pegg AE (2006) Regulation of ornithine decarboxylase. Journal of Biological Chemistry 281(21):14529–14532.

48. Almrud JJ, et al. (2000) Crystal structure of human ornithine decarboxylase at 2.1 Å resolution: structural insights to antizyme binding. Journal of molecular biology 295(1):7–16.

49. D’Assoro AB, Haddad T, & Galanis E (2016) Aurora-A kinase as a promising therapeutic target in cancer. Frontiers in oncology 5:295.

50. Alao JP (2007) The regulation of cyclin D1 degradation: roles in cancer development and the potential for therapeutic invention. Molecular cancer 6(1):1–16.

51. Daniel J, Coulter J, Woo J-H, Wilsbach K, & Gabrielson E (2011) High levels of the Mps1 checkpoint protein are protective of aneuploidy in breast cancer cells. Proceedings of the National Academy of Sciences 108(13):5384–5389.

52. Chen X, Liu W, & Liu B (2021) Ginsenoside Rh7 suppresses proliferation, migration and invasion of NSCLC Cells through targeting ILF3-AS1 mediated miR-212/SMAD1 axis. Frontiers in Oncology 11:1326.

53. Yokomizo A, et al. (1999) Overexpression of the wild type p73 gene in human bladder cancer. Oncogene 18(8):1629–1633.

54. Sanders AJ, Martin TA, Ye L, Mason MD, & Jiang WG (2011) EPLIN is a negative regulator of prostate cancer growth and invasion. The Journal of urology 186(1):295–301.

55. Jiang WG, et al. (2008) Eplin-alpha expression in human breast cancer, the impact on cellular migration and clinical outcome. Molecular cancer 7(1):1–10.

56. Subramanian D, Bunjobpol W, & Sabapathy K (2015) Interplay between TAp73 protein and selected activator protein-1 (AP-1) family members promotes AP-1 target gene activation and cellular growth. Journal of Biological Chemistry 290(30):18636–18649.

57. Li D, Jackson RA, Yusoff P, & Guy GR (2010) Direct association of Sprouty-related protein with an EVH1 domain (SPRED) 1 or SPRED2 with DYRK1A modifies substrate/kinase interactions. Journal of Biological Chemistry 285(46):35374–35385.

58. Dulloo I & Sabapathy K (2005) Transactivation-dependent and-independent regulation of p73 stability. Journal of Biological Chemistry 280(31):28203–28214.

59. Shevchenko A, Tomas H, Havli J, Olsen JV, & Mann M (2006) In-gel digestion for mass spectrometric characterization of proteins and proteomes. Nature protocols 1(6):2856–2860.

60. Sanjana NE, Shalem O, & Zhang F (2014) Improved vectors and genome-wide libraries for CRISPR screening. Nature methods 11(8):783–784.

61. Zhao Y, et al. (2011) Analgesic‐antitumor peptide inhibits proliferation and migration of SHG‐44 human malignant glioma cells. Journal of cellular biochemistry 112(9):2424–2434.

